# Measles fusion complexes from central nervous system clinical isolates: decreased interaction between hemagglutinin and fusion proteins

**DOI:** 10.1101/2021.06.18.449082

**Authors:** Cyrille Mathieu, Tiago Nascimento Figueira, Amanda R. Decker, Marion Ferren, Tiziana F. Bovier, Eric M. Jurgens, Tara C. Marcink, Anne Moscona, Matteo Porotto

## Abstract

Measles virus (MeV) viral entry is mediated by a fusion complex comprised of a receptor binding protein (hemagglutinin, H) and a fusion protein (F). The wild type H/F complex requires interaction with specific proteinaceous receptors (CD150/SLAM and nectin-4) in order to be activated. In contrast the H/F complexes isolated from viruses infecting the central nervous system (CNS) do not require a specific receptor. A single amino acid change in the F protein (L454W) was previously identified in two patients with lethal sequelae of MeV CNS infection, and the F bearing this mutation mediates fusion even without the H protein. We show here that viruses bearing the L454W fusion complex are less efficient than wt virus at targeting receptor expressing cells and that this defect is associated with a decreased interaction between the H and the F proteins.

**Importance:** Measles (Mev) infection can cause serious complications including measles inclusion body encephalitis (MIBE) and subacute sclerosing panencephalitis (SSPE). MIBE and SSPE are relatively rare but lethal. We have shown that the fusion complex of CNS adapted clinical samples can spread in the absence of known receptor. We now provide evidence that HRC mutations leading to CNS adaptation come at a cost to the efficiency of viral entry.

**One Sentence Summary:** Measles CNS adapted fusion complexes have altered H/F interaction.

## Introduction

Despite the availability of an effective measles virus (MeV) vaccine and efforts to increase vaccine coverage by the WHO, UNICEF, and their partners, MeV has not been eradicated and the estimated global measles death toll rose from 89,780 in 2016 to 207,500 in 2019(1). The SARS-CoV-2 pandemic and associated countermeasures reduced the incidence of other respiratory viral infections but also halted vaccination, increasing the risk of measles outbreaks in the future(1).

MeV is the most infectious virus known to humans, transmits via the air, and causes systemic infection. The virus enters cells via two cellular receptors: CD150/SLAM expressed by subsets of immune cells(2) and nectin-4 expressed by epithelial cells(3, 4). Central nervous system (CNS) sequelae associated with active viral infection include subacute sclerosing panencephalitis (SSPE) and measles inclusion body encephalitis (MIBE) (5-10). These CNS complications have been associated with viral evolution and specific CNS adaptation, specifically with alterations in the viral fusion mechanism(11, 12).

MeV expresses two envelope glycoproteins that make up the fusion complex. The H protein mediates receptor binding, followed by triggering of the F protein, which leads to merger of the viral envelope with target cell membranes. We refer to the H/F pairs of MeV as the viral fusion complex, since these proteins act in concert. F is synthesized as a precursor (**F**_**0**_) that is cleaved within the cell to yield the pre-fusion F complex comprising three C-terminal **F**_**1**_ subunits associated non-covalently with three N-terminal **F**_**2**_ subunits. This trimeric F structure is kinetically trapped in a metastable conformation, primed for fusion activation upon engagement of the H glycoprotein by the cell-surface receptor (either CD150/SLAM or nectin-4)(2-4, 13-15). After receptor engagement by H, the pre-fusion F undergoes a structural transition, extending and inserting its hydrophobic “fusion peptide” into the target cell. During entry, F refolds into a so-called “trimer of hairpins” (or 6-helix bundle) post-fusion structure that brings together the N-terminal heptad repeat (HRN) and the C-terminal heptad repeat (HRC), and the viral and cell membranes fuse(16-23). Peptides derived from the HRC region of the **F**_**2**_ ectodomain inhibit paramyxovirus entry with varying activity(19, 24-32). We and others have shown that fusion complexes from CNS isolated clinical samples are dysregulated and do not require H-receptor interaction for mediating fusion(8, 10, 33, 34), and for a specific mutated F (L454W) even the presence of the H was dispensable for cell-to-cell fusion in transfected cells(8, 10, 34). Here we investigated whether this phenotype was correlated with dimished H-F interaction and we highlight that fusion in the absence of receptor interaction is the combined effect of a decrease in the stability of F as well as a decrease in F’s interaction with H.

## Results

### Measles virus F glycoprotein from neuropathogenic viruses

MeV F glycoproteins with specific single residue alterations (S262R, L454W, T461I, and N462K) have been associated with neuropathogenic measles strains that were either isolated from patients or generated in laboratory settings (9, 12, 34, 35). We mapped these mutations onto x-ray structures of the pre-fusion (36) conformations of F (Fig.1A), highlighting the locations of these mutations in MeV F. The S262R mutation occurs at the interface of three protomers in the head region of the pre-fusion structure. The mutation from Ser to Arg at residue 262 results in a clash within each monomer in the existing crystal structure . These microenvironmental changes are likely to affect the conformational stability of the F. The other three mutations (L454W, T461I, and N462K) were located within the C-terminal heptad repeat domain (HRC). In the pre-fusion structure, the L454W mutation would also perturbs the current pre-fusion structure. The T461I and N462K mutations occur in a well-ordered α-helical region of the HRC domain. These three mutations (L454W, T461I, and N462K) occur in the portion of the HRC domain where the head and stalk regions of the pre-fusion conformation meet. Interactions at this region may be important for stabilizing the pre-fusion state and mutations in this junction would affect stability of the MeV-F prefusion structure, as our previous data have shown (12, 34).

**Figure 1.**
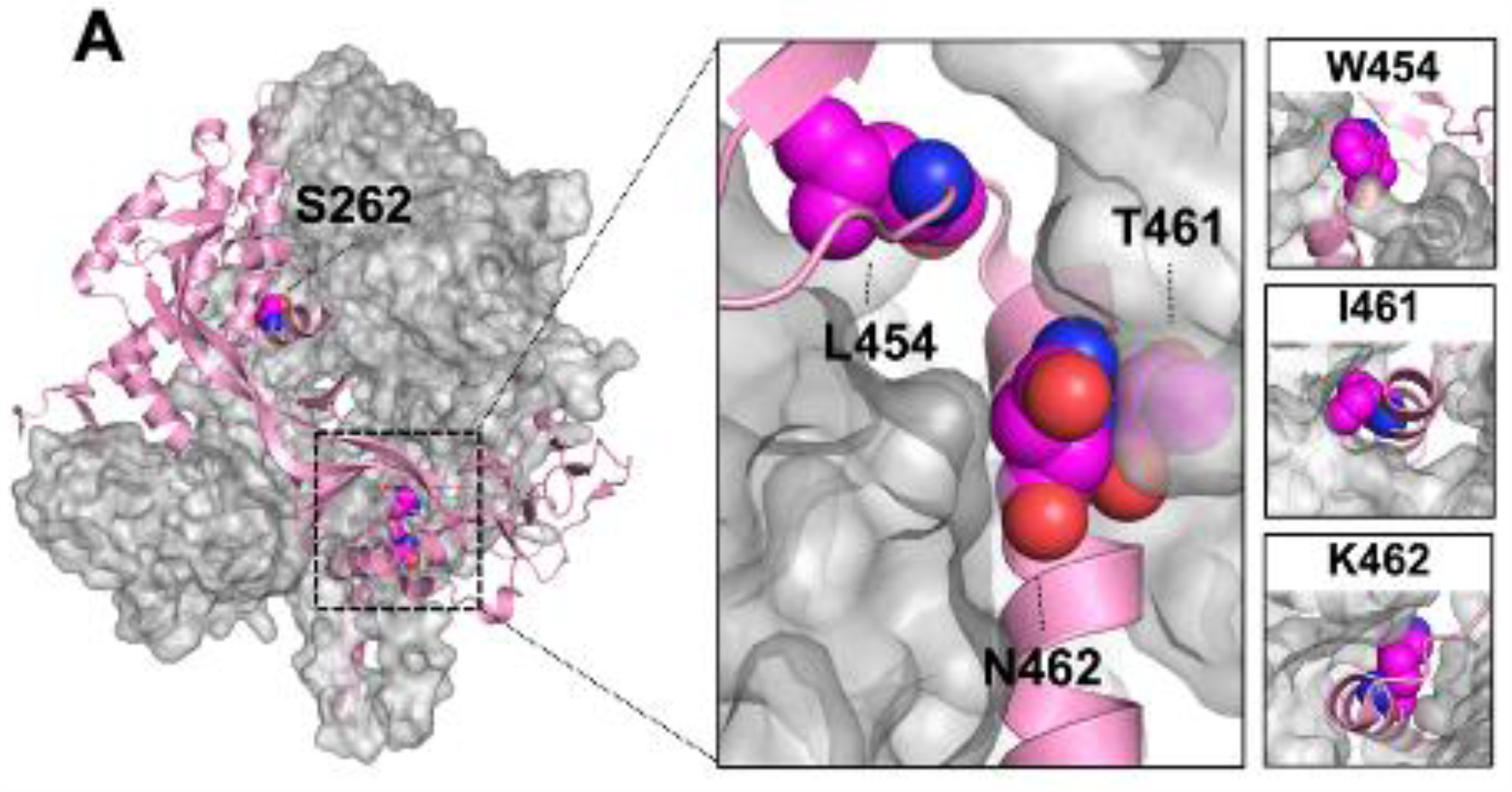
Crystal structure of the trimeric measles F (PDBID:5YXW) in the prefusion state. Monomer is shown in pink. Zoomed in view shows close proximity of residues L454, T461, and N462 near the top of the prefusion F stalk. Left-most panels show potential orientations of mutations W454, I461, and K462 as modeled by pymol with minimal sculpting to reduce steric clashes. Three substitutions (L454W, T461I and N462K) in the HRC domain in neuropathogenic strains are shown. The residue S262 (mutated here to R) was previously described in the hyperfusogenic F variant.

### Impact of hyperfusogenic mutations in F on H-F interaction

We previously suggested that the ability of HRC mutants to fuse in the absence of known receptors could be at least partly due to reduced interaction between H and F(34). The interaction between F and H and the impact of mutations in the HRC domain (*i*.*e*., L454W, T461I and N462 may be affected by thermal instability – obscuring the effect of the specific mutations --we included the highly unstable mutant F-S262R (11, 12, 33, 35) which interacts efficiently with H. H-F interaction was assessed with both the cleavable F proteins (Fig. 2A) and the cleavage site mutant (CSM) version of these F proteins (Fig. 2B), to distinguish between effects on the **F**_**0**_ and the **F**_**1**_ form of F. Results are presented as the average of 3 separate experiments ± SD (Fig.2C). Densitometric analysis was used to normalize the results to total protein content and to convert the data to graphs (average of 3 separate experiments ± standard deviation; Fig. 2C). There is significant reduction in the amount of **F**_**1**_ co-immunoprecipitated with H for the HRC hyperfusogenic mutants (*i*.*e*., L454W, T461I and N462K) compared to the *wt* F (*P* = 0.0023, *P* = 0.0030 and *P* = 0.0014, respectively; two-way ANOVA with multiple comparisons against *wt* F, corrected with the Dunnett hypothesis test). For the other hyperfusogenic mutant, F S262R, there was more co-immunoprecipitated **F**_**1**_ protein when compared to the HRC mutants. There were no differences in co-immunoprecipitation of **F**_**0**_. Thus, mutations in HRC associated with a hyperfusogenic phenotype decrease the ability of H to interact with the active **F**_**1**_ form of F.

**Figure 2.**
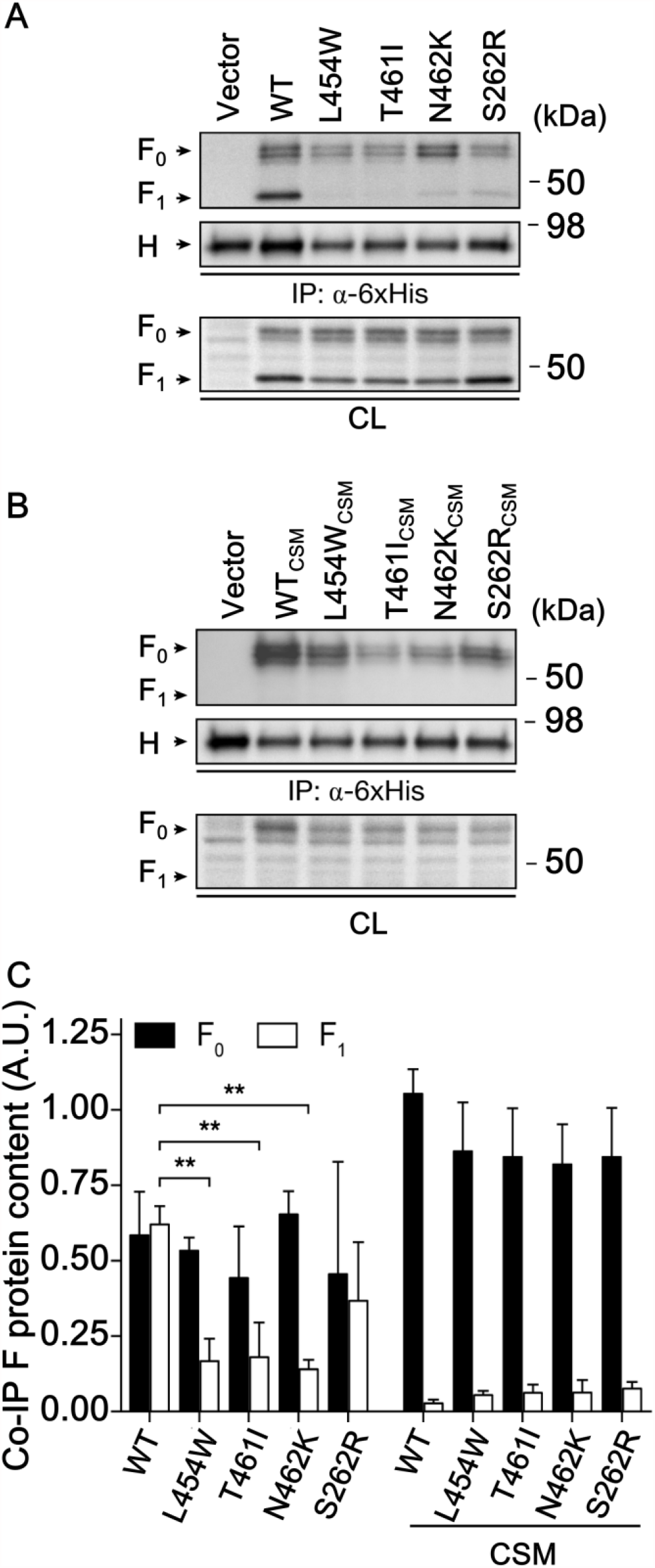
The effect of hyperfusogenic MeV F HRC domain mutations on MeV H functional and physical interactions with F_0_ and F_1_. **(A-B)** Cell lysates from 293T cell cultures co-expressing MeV H-6xHis and F *wt*, L454W, T461I, N462K or S262R proteins were immunoprecipitated with an anti-6xHis antibody. Cleavable (A) and non-cleavable (B) forms of F were included in the experiment. Precipitates were analyzed by western blotting using anti-MeV F HRC and anti-6xHis (top panels) while cell lysates were blotted using anti-MeV F HRC (bottom panels). **(C)** Densitometry measurements of co-immunoprecipitated MeV F protein detected through western blot analysis. Protein content was normalized for the total F protein in cell lysates and the immunoprecipitated H protein. Results are the average of three independent replicates. Error bars correspond to the standard deviation. **, *P* ≤ 0.01.

### Viral evolution leads to compensatory mutation

MeV bearing the L454W F protein, when grown at 37°C in cell culture, acquired a compensatory mutation (E455G) that re-balanced F’s stability and dependence on H and cellular receptor for mediating fusion (12). The E455G mutation is shown in Fig. 3A, alone and in combination with L454W. Tryptophan (W454) contains a bulky aromatic sidechain when compared to leucine (L454), while glutamic acid (E455) is larger than glycine (G455). We determined whether the E455G mutation restored H-F interaction (Fig. 3). H-F interaction for wt, L454W, E455G, and L454W/E455G F proteins is shown in Fig. 3 . Densitometric analysis was used to normalize the results to total protein content and to convert the data to graphs (average of 3 separate experiments ± SD; Fig. 3C). The *wt* versus the L454W F was significantly different (as seen in Fig.2), however the mutant E455G F and the double mutant L454W/E455G F co-immunoprecipitated similarly to the *wt* F. The E455G compensatory mutation in the HRC domain of the F resulted in a *wt* phenotype. We previously showed that E455G restored F’s stability (as measured by sensitivity to thermal activation) and restored F’s dependence on H-receptor interaction for fusion(12). Fig. 3 reveals that E455G also reestablished interaction of H with the active **F**_**1**_ form of F.

**Figure 3.**
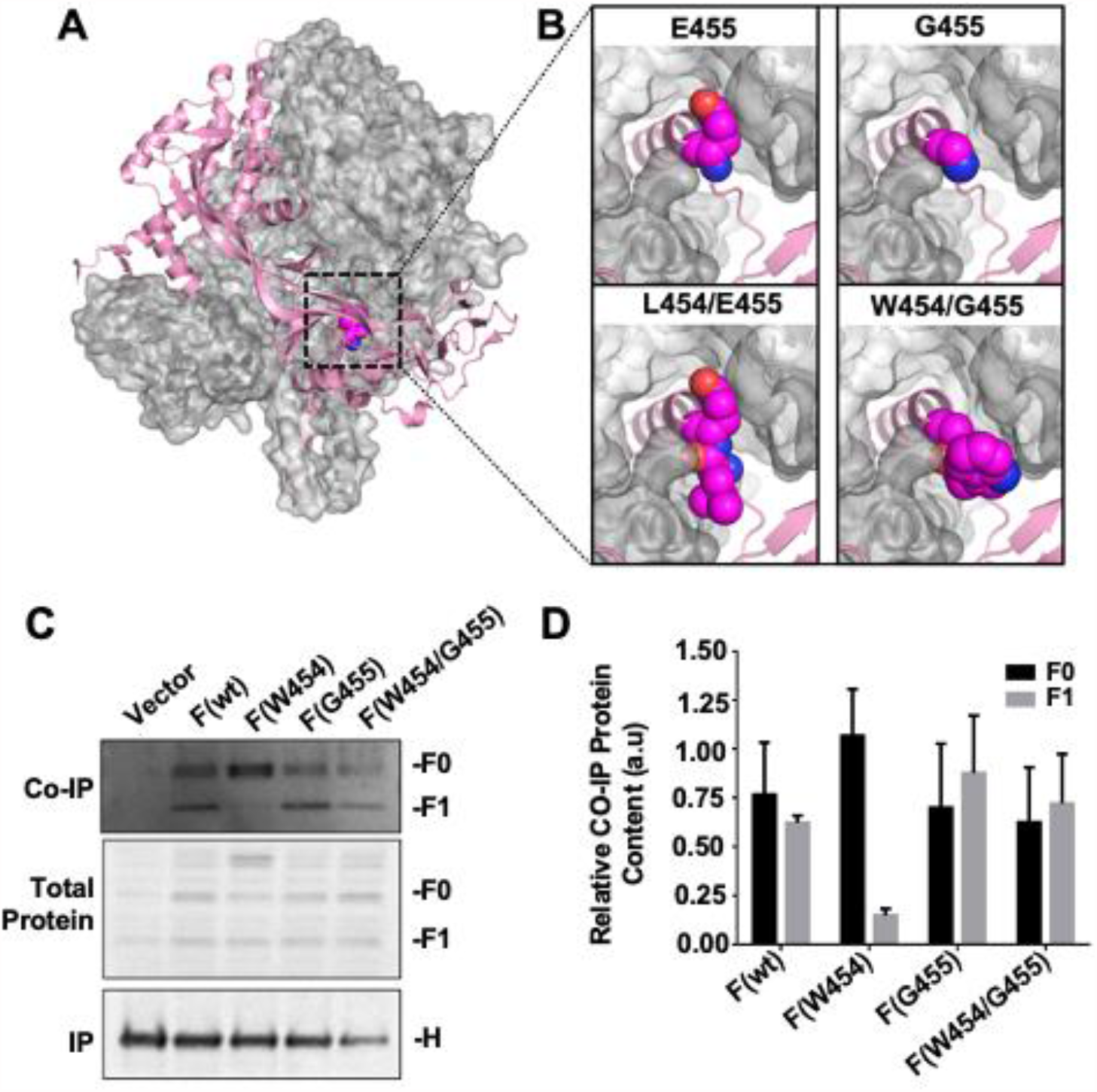
The effect of F protein’s E455G compensatory mutation on H protein’s interaction with F_0_ and F_1_. **(A)** Prefusion F crystal structure (PDBID: 5YXW). (B) Zoomed in views near the top of the prefusion F stalk region with E455 (top left) and L454/E455 (bottom left) shown as spheres. Mutations G455 (top right) and W454/G455 (bottom right) were substituted in pymol with minimal sculpting to reduce steric clashes. **(C and D)** Cell lysates from 293T cell cultures co-expressing MeV H-6xHis and F *wt*, L454W, E455G, or L454W/E455G proteins were immunoprecipitated with an anti-6xHis antibody. Cleavable (B) and non-cleavable (C) forms of F were included in the experiment. Precipitates were analyzed through western blotting using anti-MeV F HRC and anti-6xHis (top panels) while cell lysates were blotted using anti-MeV F HRC (bottom panels). **(D)** Densitometry measurements of co-immunoprecipitated MeV F protein detected by western blot analysis. Protein content was normalized for the total F protein in cell lysates and the immunoprecipitated H protein. There was no statistical difference (unpaired T-test) between the levels of **F**_**0**_ that were co-immunoprecipitated for each allele. There was also no statistical difference between F(*w*t) and either allele that contained E455G (single and double mutant) for **F**_**1**_ levels. However, the difference between the F(*wt*) and F(L454W) **F**_**1**_ that co-immunoprecipitated with H was extremely statistically significant (P = 0.0007). Error equals +/- SEM. N = 3 biological replicates

### Loss of H-F interaction delays viral entry

One of the consequences of the loss of interaction between F and H is the loss of protection of F from spontaneous triggering(37). Additionally, hyperfusogenic MeV F HRC mutants proteins are less stable in their pre-fusion state, as measured by their temperature sensitivity, compared to the wt F protein. The hyperfusogenic F proteins mediate fusion even in the absence of either CD150/SLAM or nectin-4 receptors(10, 34). We have shown that this decreased F stability, however, results in viruses that are inactivated at lower temperatures than wt virus (10), because F transitions to its post-fusion state more readily, rendering the viral particle non-infectious. We asked whether the destabilizing mutations in F of the neuropathogenic variants affect viral entry into SLAM-expressing cells (38). To do so, we used an assay developed for a related paramyxovirus, human parainfluenza virus 3 (HPIV3). For HPIV3 we have shown that the kinetics of F protein activation and viral entry modulate the potency of HR derived peptides(37, 39), and therefore this modulation can be used as a tool to quantitate F activation and entry. Applying this strategy to MeV, we determined the sensitivity of MeV F to inhibition by MeV HR derived peptides, using a previously described dimeric HR lipopeptide, HRC4. In Fig. 4A, 1µM of HRC4 peptide was added at time points from 0 to 2 hours during infection of Vero-SLAM cells. Peptide addition at time zero inhibited 100% of infection for all the viruses (wt and the three viruses bearing the mutated F proteins). Peptides added after 1 hour decreased inhibition to 30% for wt virus, meaning that most of the wt virus had already entered, while the mutants bearing F T461I and N462K were 50% inhibited and the mutant bearing F L454W was 70% inhibited. After 90 min, the peptide continued to inhibit the mutants (from 20 to 30% of viral entry). No inhibitory effect was observed after 2 hours. This experiment reveals that for wt viruses, the window of HRC4 inhibitor sensitivity is significantly shorter than that of the viruses bearing the mutated Fs. The longer time of inhibitor susceptibility of the HRC mutant Fs suggests that wt H-F is more efficient for entry in the presence of receptors.

**Figure 4.**
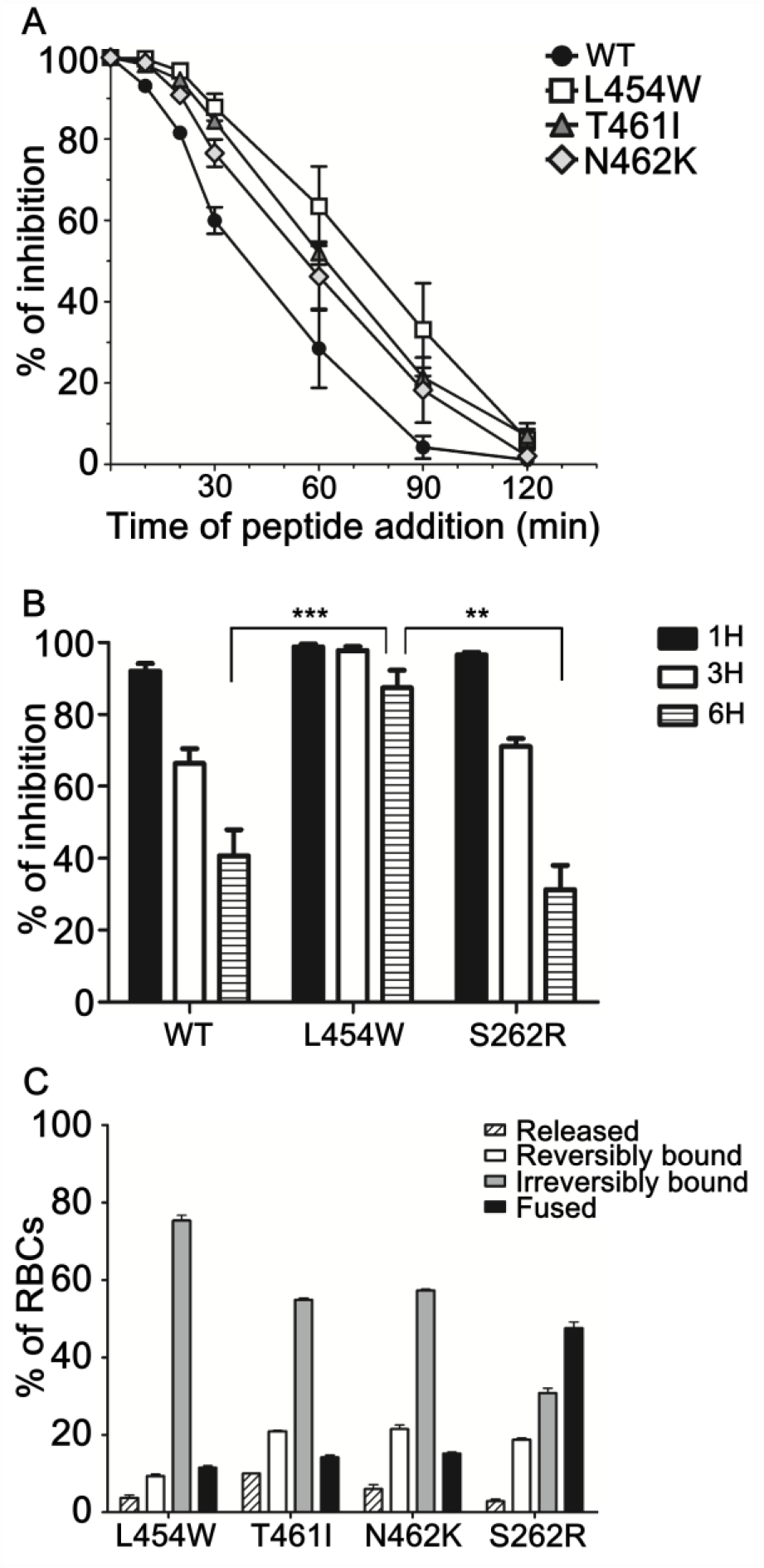
HRC hyperfusogenic mutants fuse less in presence of entry receptor. (**A**) Inhibition by MeV HRC4 peptides added at different times after infection reveals differences in the rate of F activation. Vero-SLAM cells were infected with wt or the indicated viruses at a multiplicity of infection of 6.7 × 10^−4^. The MeV HRC4 fusion inhibitory peptide was added at the time points noted to a final concentration of 1 μM. Cells were overlaid with agarose at 90 min and plaques were stained and counted at 36 h. The percent inhibition of viral entry, normalized to 100% inhibition at time zero, is shown as a function of the time of peptide addition. Data points are means (± standard error) for triplicate experiments. **-** (**B**) Fusion of MeV H/F co-expressing cells with SLAM-bearing cells in the presence of unconjugated peptides (HRC1 at 1 μM) was quantitated at 1 h, 3h or 6 h using a β-galactosidase complementation assay. Results are presented as percent reduction in luminescence (*y* axis) compared with no treatment. Each point is the mean (± standard error) of results from 3 to 5 separate experiments (**, *P* < 0.01; ***, *P* < 0.001 [Mann-Whitney-U test]). (**C**) Role of receptor binding protein (H-HN) after F has inserted its hydrophobic fusion peptide into the target cell: progression to membrane merger in the presence of H-F interaction. If H-F interaction is functional, processed F is activated to the stage of fusion peptide insertion captured by unconjugated peptides, and then proceeds to membrane merger. Monolayers of cells co-expressing H-HN and processed F proteins (as indicated on the axis) were allowed to bind to receptor-bearing RBCs at 4°C. Unbound RBCs were washed away, and cells were incubated with standard HRC1 peptides (1 μM) at 37°C for 60 min. The values on the *y* axis reflect the quantity of RBCs that were released by H-HN neuraminidase (striped bar), reversibly bound by H-HN-receptor interaction (white bar), irreversibly bound (gray bar), or fused (black bar). The values are means (±standard deviations) of results from triplicate samples and are representative of the experiment repeated at least four times.

### Easier triggering of HRC hyperfusogenic mutants correlates with longer time to completion of fusion

For HPIV3, fusion requires engagement of the receptor binding protein (the hemagglutinin neuraminidase, HN) beyond initial triggering of the F protein. This engagement of HN is essential for F’s function until membrane merger (37). A similar mechanism of ongoing activation of the fusion process by receptor-engaged H could explain our observations for MeV. We showed that constant interaction with F by receptor-engaged H may allow wt H-F to fuse even in the presence of F-targeted anti-MeV peptides (40). As above, we also previously showed that for HRC peptide fusion inhibitors, adding a lipid moiety improves antiviral potency over time(40). In this case a cholesterol-conjugated dimeric peptide (HRC4) was inhibitory as long as 6h after *in vitro* (41). Without a lipid moiety (HRC1), peptide was effective only at early time points after infection.

We hypothesize that the loss of of H-F interaction affects both the triggering of HRC mutants and their ability to fuse in the presence of inhibitor. In Fig. 4B we assessed weak peptide inhibitor (HRC1) for inhibition of wt and mutant F fusion. The assay measures the fusion of cells that express viral envelope glycoproteins (MeV IC323 H/F) with cells that express the MeV receptor SLAM. HRC1 peptide inhibited fusion at early time points (1h, Fig. 4 B). As expected, this inhibition decreased below 70% after 3h and below 40% for the wt F and S262R F respectively after 6h. In contrast, for the H/L454W F fusion complex, HRC1 peptide inhibited over the time course of this fusion assay, significantly better than for the wt complex (**p value, Mann-Whitney U-Test). A parallel assay was performed with the HRC4 peptide, which completely inhibited fusion of all MeV Fs (Fig. S1).

HRC peptides are thought to stabilize F protein intermediates during the fusion process after insertion into the target cell membrane but before refolding to the post-fusion state. The stabilization of the transient intermediate state of F by HRC peptides was used to identify discrete steps in the fusion process (37, 42). In Fig. 4C we determined whether the HRC1 fusion blockade for the HRC mutated F occurs at the same transitional state. Red blood cells (RBCs) were allowed to interact with cells expressing the MeV F hyperfusogenic mutants along with a chimeric receptor binding protein expressing the MeV H stalk and the globular head of HPIV3 HN. This chimeric protein can engage the sialic acid receptor on the RBCs and initiate MeV F-activation for the neuropathogenic F proteins (8, 34, 43). The read-out for peptide inhibition is irreversible binding of the RBCs to the F-expressing cells and/or fusion. This assay provides a measure of the efficiency with which HRC1 binds and retains F in its transitional intermediate state, preventing membrane merger (Fig.4C). 293T cells co-expressing the chimeric receptor binding protein MeV H-HN and MeV F (hyperfusogenic mutants) were incubated with RBCs at 4 °C, then transferred to 37 °C in the presence of HRC1 peptides (1μM). Zanamivir, a small molecule sialic acid analog that blocks HN-receptor interaction was added after 1 h to disengage cells that were bound by HN-globular head engagement alone. Note that wt F was not used in this assay since it is too stable to be activated under these conditions. The proportion of RBCs either free, reversibly bound by HN-receptor interaction (and therefore released by zanamivir), irreversibly bound by F protein bridging (following insertion of the fusion peptide in the target cell membrane), or fused was quantified. For all three MeV F HRC mutants the larger majority of the cell population remained irreversibly bound and very little fusion was observed, confirming the ability of HRC1 to block F in its transitional state after insertion of the fusion peptide in the target cell membrane. In contrast, the percentage of irreversibly bound cells decreased to 30% concomitantly with an increase in fusion up to 50% for the S262R hyperfusogenic mutant, confirming that this F can “override” inhibition by the peptide. The HRC1 peptide stabilizes the intermediate fusion state for all the F proteins that bear mutations in the heptad repeat domain. Taken together, these results confirm that while the reduction in H-F interaction allows HRC hyperfusogenic mutants to fuse in the absence of receptor — which seems to be a benefit in the CNS— it also decreases their fusion compared to wt virus in presence of receptor by reducing complete fusion after triggering. Outside the CNS, this reduced fusion is a disadvantage.

## Discussion

The altered HRC region of the CNS-adapted MeV F proteins produced viruses that can fuse and infect without known receptors for H. While H-F interaction is essential for fusion and infection by wt MeV, these F proteins (L454W, T461I and N462K; fig 1A) interact less with H but nevertheless mediate infection *in vitro* (L454W, T461I and N462K) (10, 34), in models of CNS infection (L454W, and T461I)(12) and *in vivo* (L454W) (10) more effectively than wt virus in the absence of SLAM or Nectin-4 receptors. As anticipated based on previous data(10, 12, 34), the hyperfusogenicity conferred by the alterations in the MeV F HRC region affects both functional and physical interactions with H. The E455G revertant restored wt-like properties in the context of the L454W-bearing F, and when introduced singly into the F protein led to an extremely stable F(12) . We speculate that introducing W454 into the prefusion F stalk results in local destabilization that enhances F activation, even in the absence of receptor. The nearby mutation G455 may alter the structure sufficiently to restore the “wt” properties. When the viral quasispecies formed by L454W and L454W/E455G F bearing viruses is cultivated in human brain organoids, the L454W/E455G F bearing virus is eliminated within 10 days confirming that the L454W F bearing virus is fit for growth in CNS tissues(12). The virus bearing the S262R mutation in F is neuropathogenic *in vivo* (11, 35), and we observed no significant differences in H-F interaction. A S262G mutation in F (along with several other mutations) was found in virus from a SSPE clinical case (8). Thus, while destabilizing the F protein significantly improves MeV’s ability to spread in the brain, the loss of H-F interaction may not be necessary for CNS adaptation.

MeV H-F interaction is key for the steps after F insertion in the target membrane, to complete fusion. The viruses bearing F mutations that lead to CNS spread are more susceptible to fusion inhibitory peptides(10). We attribute this, at least in part, to the fact that the H does not continue to activate F after initial triggering.

In Fig. 5 we propose models of viral infection with wt MeV and the CNS adapted variants, incorporating the findings in this work as well as work from others. A common pattern for neuropathogenic F variants is decreased stability of F (as assessed by sensitivitiy to heat) and a fusion complex that mediates cell-to-cell fusion in the absence of either CD150/SLAM or nectin-4(8, 10, 12, 34, 44). These properties confer an infectivity cost, since they are significantly slower to complete the fusion process after inital triggering. These features together with lower viability (10) make these viruses adapt to cell-to cell transmission after initial infection but also likely mean that they are less transmissible.

**Figure 5.**
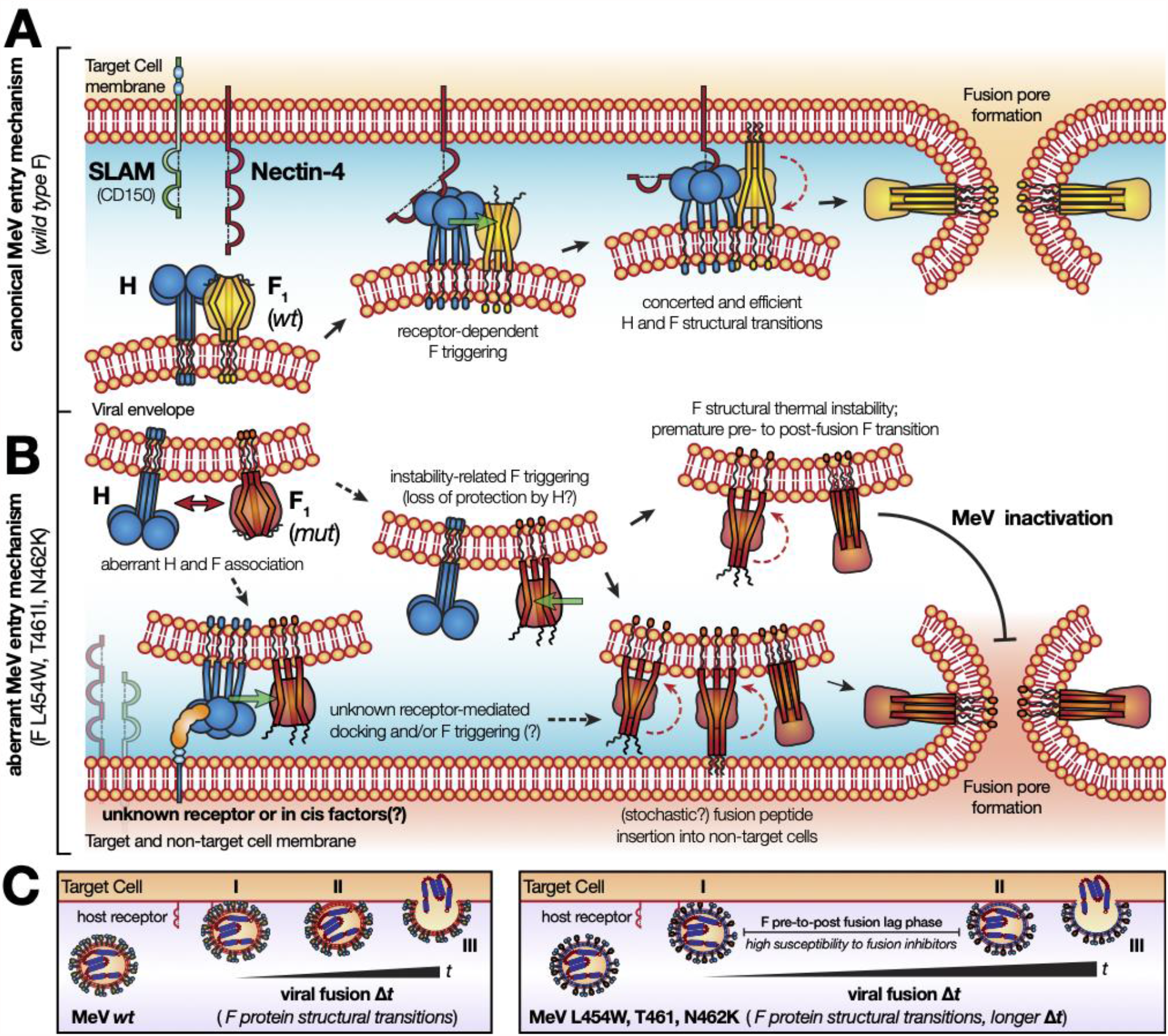
Model for the hyperfusogenic entry mechanism of neuropathogenic MeV mutant strains. **(A)** The *wt* MeV entry mechanism is characterized by the concerted action of the H and F glycoprotein complex. Upon H-mediated CD150/SLAM or Nectin-4 receptor recognition, and subsequent F triggering, both H and F undergo a series of conformational changes that guide fusion of the cell membrane and viral envelope. H anchors the virus to the cell surface and activates the F to insert its fusion peptide trimers into the cell membrane and undergo a conformational transition, the driving force for lipid mixing and fusion. **(B)** The hyperfusogenic phenotype of neuropathogenic MeV variants is associated with single amino acid mutations in F (L454W, T461I and N462K). Altered association with H triggers the mutated F into unstable fusogenic conformations. The lower energy barrier between the pre- and post-fusion (6HB) conformations leads to premature structural transitions of F and consequent viral attenuation/inactivation. Yet, the mutated F can insert into non-targeted cell types, such as neurons, promoting MeV aberrant infection. **(C)** The coordinated structural transitions of H and F promote entry by *wt* MeV after receptor engagement (left panel). In hyperfusogenic MeV variants, with loss of H-to-F interaction, entry kinetics are slower, with a lag between the F pre- and post-fusion conformations (right panel). This feature enhanges susceptibility to F-specific fusion inhibitory peptides acting at this stage.

## Figure legends

**FIG S1.**
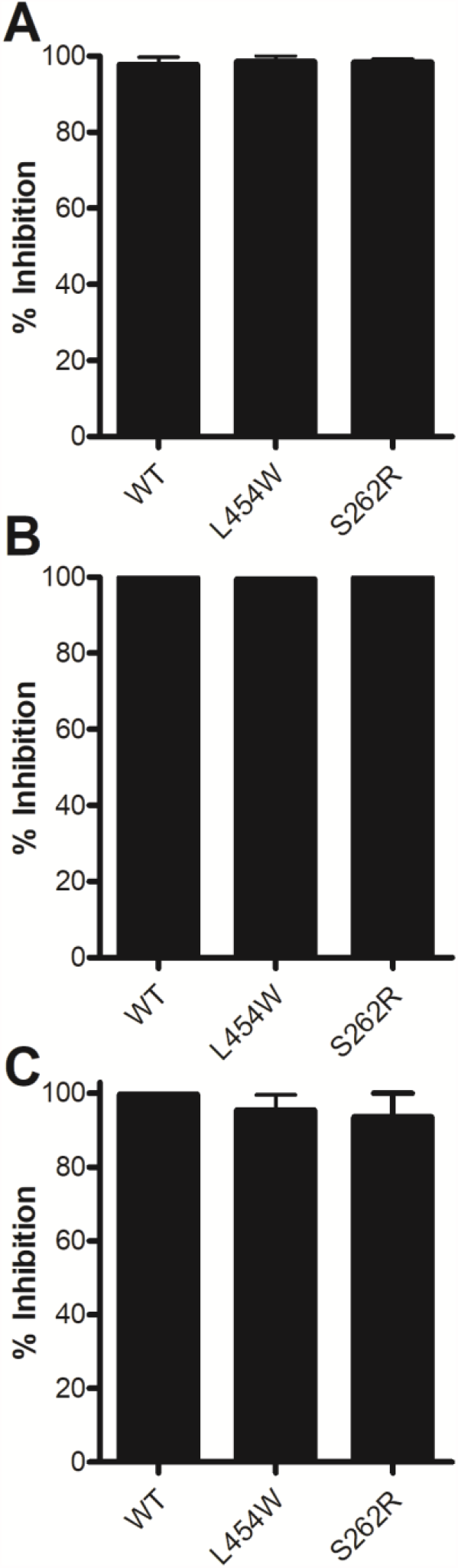
Fusion of MeV H/F co-expressing cells with SLAM-bearing cells in the presence of unconjugated peptides (HRC4, dimer peptide with cholesterol, 1 μM) was quantitated at 1 h (**A**) 3h (**B**) or 6 h (**C**), using a β-galactosidase complementation assay. Results are presented as percent reduction in luminescence (*y* axis) compared with no treatment. Each point is the mean (± standard error) of results from 3 to 5 experiments.

## Materials and Methods

### Peptides

MeV F derived fusion inhibitor peptides HRC1 and HRC4 were previously described (40). Briefly 36aa long peptides derived from the heptad repeat region at the C-terminal of the MeV F protein were synthesized (using the wt sequence or the L454W sequence). Monomeric unconjugated (HRC1) or dimeric cholesterol conjugated (HRC4) forms of the peptides were used in this study.

### Plasmids and reagent

The genes of MeV IC323 H and F proteins were codon optimized, synthesized, and sub cloned into the mammalian expression vector pCAGGS. Plasmids encoding nectin 4 and CD150/SLAM were commercially acquired.

### Cells

293T (human kidney epithelial) and Vero-SLAM (African green monkey kidney) cells were grown in Dulbecco’s modified Eagle’s medium (DMEM; Gibco, Invitrogen) supplemented with 10% fetal bovine serum (FBS) and antibiotics in 5% CO_2_. The Vero-SLAM culture medium was supplemented with Geneticin.

### HRC4 Peptide time addition related inhibition

Dulbecco’s Modified Eagle’s medium (DMEM) containing the indicated viruses (MeV IC323-EGFP, MeV IC323-EGFP-F L454W, MeV IC323-EGFP-F T461I, MeV IC323-EGFP-F N462K all these viruses are from(34)) was used to infect sub-confluent VERO-SLAM cells in 6 well plates (100pfu/well) for 2h at 32°C. MeV HRC4 dimeric fusion inhibitory peptide (1μM) was added to the medium at time points from 15 to 120 min after the beginning of infection. After 2h of incubation with the virus, medium was replaced with medium containing Avicel. Viral titers were assessed after 3 days of incubation at 32°C by immune staining.

### Beta-Galactosidase (Beta-gal) complementation-based fusion assay

The beta-gal complementation-based fusion assay was performed as described previously(34). Briefly, 293T cells transiently transfected with the constructs indicated above and the omega reporter subunit were incubated for the indicated period with cells coexpressing viral glycoproteins and the alpha reporter subunit in presence or not of MeV F HRC derived peptide (40).

### RBC fusion assay

RBC fusion assays were performed on HEK 293T cells transiently expressing MV H_Y17H HPIV3_T193A chimerae (43) and the indicated MeV Fs. Cell monolayers were washed three times with serum-free medium, placed at 4°C with 1% RBC in DMEM for 30 min, then treated with the indicated peptide and placed at 37°C. Zanamivir was added at a final concentration of 10 mM and the cells were incubated at 4°C, rocked, and the liquid phase was collected in V-bottomed 96-well plates for measurement of reversibly bound RBCs. The cells were then incubated at 4°C with ACK-Lysing buffer, and the liquid phase was collected in V-bottomed 96-well plates for measurement of irreversibly bound RBCs. The cells were then lysed in lysis buffer and transferred to flat-bottom 96 well plates for quantification of fused RBCs. The amount of RBCs in each of the four compartments (released, reversibly bound, irreversibly bound, and fused) was determined by measurement of absorption at 410 nm.

### Co-Immunoprecipitation

293T cells were seeded in biocoated 6-well plates (Corning) at 5×10^5^ cells/well and incubated for 24h, at 37° C. Cells were transfected with plasmid vectors encoding for MeV H-6xHis, MeV F (*wt*, L454W, T461I, N462K,E455G, L454W/E455G and S262R) and mCherry (6:4:1 mixtures; 2.2 μg/well) using Lipofectamine® 2000 (Invitrogen), according to the manufacturer’s recommendations. Transfected cultures were grown at 32° C, overnight, in the presence of HRC1 peptide (1 μM), to minimize cell-cell fusion. mCherry fluorescence was used to monitor cell transfection efficiency after 18h. Prior to cell lysis, cell protein expression was synchronized with cycloheximide (Sigma, 0.1 mg/mL), followed by membrane protein cross-linking with 3,3’-dithiobis(sulfosuccinimidyl propionate) (Sigma; 1 mM), at low temperature. The cross-linking reaction was quenched with 20 mM Tris, 150 mM NaCl, pH 7.5. Cells were lysed with 50 mM HEPES, 100 mM NaCl, 0.05 g/mL dodecyl maltoside, pH 7.5, supplemented with complete protease inhibitor cocktail (Roche). Lysates were centrifuged at 16000 *g* for 10 min to remove nuclei and cell debris, and the supernatant was collected for immunoprecipitation and total protein content analysis. MeV H-6xHis protein was immunoprecipitated from cell lysates using Dynabeads® (Thermo, 1 mg/mL) coated with a 6xHis tag-specific antibody (mouse monoclonal, Thermo, MA1-21315).

### For Fig.2

Co-immunoprecipitated and cell lysate proteins were analyzed by western blotting, using primary antibodies specific for MeV F HRC (rabbit polyclonal, Genscript, 503028-1) and 6xHis tag (rabbit polyclonal, Thermo, PA1-983B), followed by an HRP-conjugated anti-rabbit secondary antibody (Kindle Biosceiences, R1006). Western blots were developed using the SuperSignal West Femto substrate (Thermo) and imaged on a *Kwik*Quant™ Imager UV (Kindle Biosciences).

### For Fig. 3

Co-immunoprecipitated and cell lysate proteins were analyzed by western blotting, using primary antibodies specific for MeV F HRC (rabbit polyclonal, Genscript, 503028-1) and 6xHis tag (rabbit polyclonal, Thermo, PA1-983B), followed by WesternBreeze Chromogenic Immunodetection Protocol for detection.

